# Organelle-Specific pH Quantification Using a Protein-Based Quantum Sensor

**DOI:** 10.64898/2026.10.19.760020

**Authors:** Masaaki Seki, Shohei Aizawa, Yoko Shigeno, Nobuo Maita, Sisi Chen, Masato Kato, Satoshi Ogasawara, Takeshi Murata, Lewis M. Antill, Hitoshi Ishiwata

**Author notes:** first author(s).

## Abstract

Quantum sensing enables measurements of physicochemical changes within living cells at the organelle level. Conventional nanoparticle-based sensors, however, face limitations in subcellular targeting, intracellular tracking, and following dynamic biological processes across distinct intracellular environments. Here, we investigate MagLOV2, a genetically encoded quantum sensor, by examining its organelle-specific localization, magnetic field effects (MFEs) during mitophagy, and pH dependent response kinetics. MagLOV2 retained magnetic responsiveness in the cytoplasm, nucleus, and mitochondria, with fluorescence contrasts of 17.1%, 12.3%, and 28.0%, respectively. Optimization of mitochondrial targeting increased fluorescence 2.7 fold. During CCCP induced mitophagy, lysosomal colocalization of MagLOV2 increased from 20.75% to 89.73%, accompanied by slower MFE responses in acidic compartments. Purified MagLOV2 showed response times ranging from 8.27 s at pH 4.5 to 1.56 s at pH 9.5. Structural, spectral, and kinetic analyses suggested that protonation dependent flavin photochemistry and the surrounding protein environment contribute to this behavior. These measurements demonstrate that genetically targeted MagLOV2 can follow organelle transport and report local acidification through changes in magnetic-response kinetics.

## INTRODUCTION

Quantum sensing uses observables that depend on quantum states, including spin resonance, relaxation dynamics and magnetic-field-dependent optical responses, to measure local physicochemical environments. Intracellular measurements of temperature, pH, reactive oxygen species, radicals and paramagnetic ions have been achieved using methods such as optically detected magnetic resonance (ODMR) and spin relaxation measurements [1]. Most established approaches rely on externally introduced probes, including nitrogen-vacancy (NV) centres in nanodiamonds and molecular quantum nanosensors (MoQNs) [1,2]. Although these probes offer sensitive measurements in living cells, their localisation depends on delivery and intracellular trafficking rather than genetic specification. It can therefore be difficult to assign their signals to particular biological components as those components move between organelles.

Genetically encoded protein-based quantum sensors offer an alternative because they can be expressed within living cells and directed to selected subcellular compartments using targeting sequences [3]. Recent studies of MagLOV and other engineered fluorescent and flavoprotein systems have demonstrated optically detectable magnetic-field effects (MFEs) in biological environments [4–7]. In flavin-containing proteins, photoinduced electron transfer can generate spin-correlated radical pairs. Singlet–triplet interconversion in these pairs influences subsequent reaction pathways and fluorescence yields (Fig. 1b) [8–13]. Most demonstrations, however, have concentrated on fluorescence contrast under an applied magnetic field or responses to external magnetic perturbations. Whether additional magnetic-response observables can characterise intracellular environments as labelled organelles are transported remains less well explored.

**Fig. 1.**
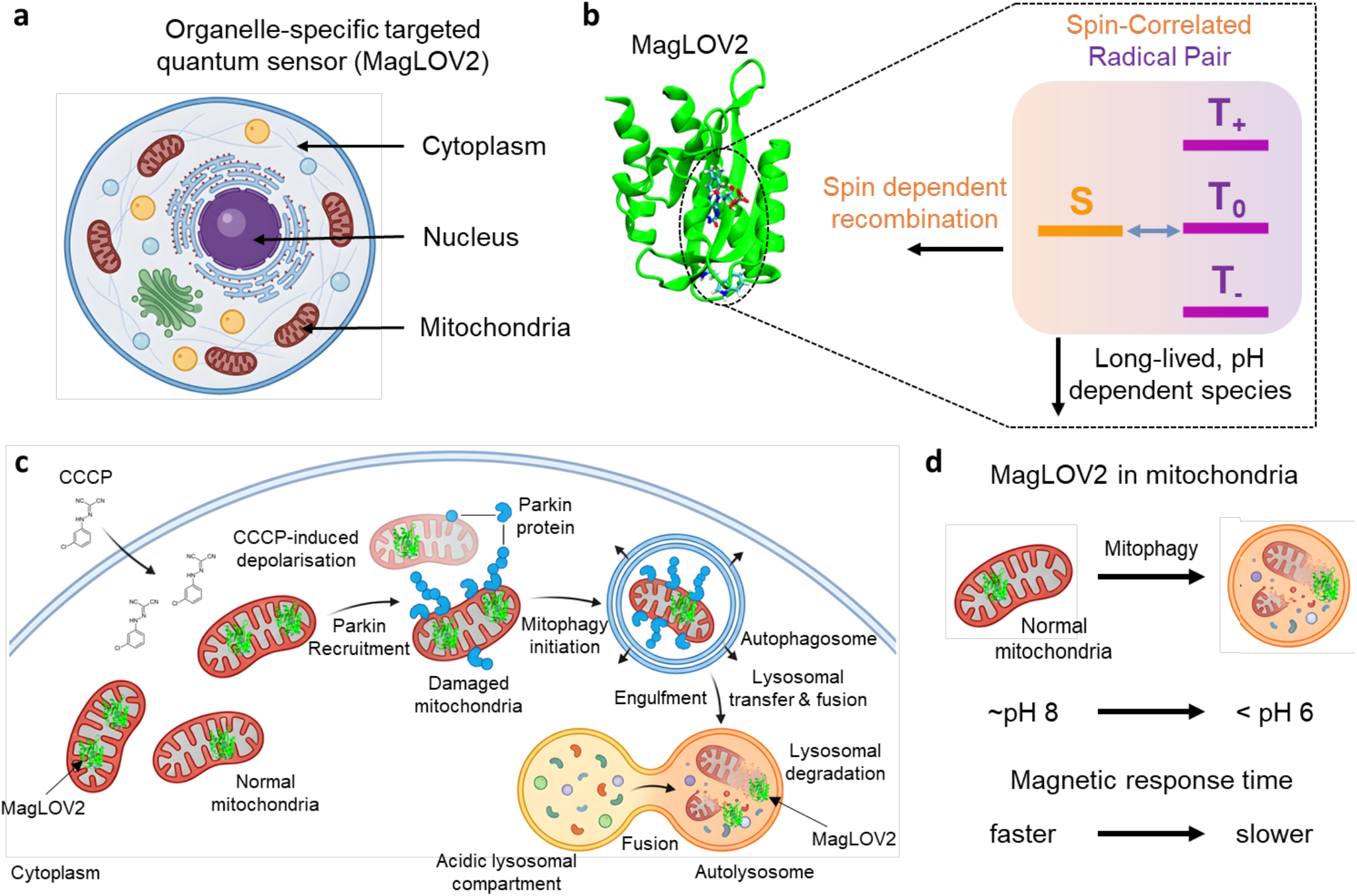
Schematic illustration of genetically encoded MagLOV2 quantum sensing and its application to mitophagy. **a**, Genetic expression and subcellular targeting of MagLOV2 in living mammalian cells, enabling localization to the cytoplasm, nucleus, and mitochondria. **b**, Proposed magnetic-field-sensitive mechanism of MagLOV2. Photoinduced electron transfer between flavin mononucleotide (FMN) and tryptophan W89 generates a spin-correlated radical pair, whose singlet (S) and triplet (T_+_, T_0_, T_–_) spin states undergo magnetic-field-dependent spin evolution. Spin-dependent recombination influences the fluorescence response, providing an optical readout of radical-pair dynamics. **c**, Schematic of CCCP-induced mitophagy in Parkin-expressing cells. Mitochondrial depolarization promotes Parkin recruitment, followed by the engulfment of damaged mitochondria by autophagosomes and their subsequent delivery to lysosomes for degradation. Mitochondrially targeted MagLOV2 accompanies the mitochondrial material during this process. **d**, Illustration of MagLOV2-labelled mitochondria before and after lysosomal delivery. The transition from the normal mitochondrial environment to the acidic lysosomal compartment alters the local physicochemical conditions surrounding MagLOV2, producing a slower magnetic-field response consistent with pH-dependent radical-pair photochemistry.

The magnetic response of a radical-pair system depends on the applied field and on the physicochemical conditions that regulate radical-pair formation, spin evolution and recombination [14]. MFEs in molecular and protein-based radical pairs are sensitive to temperature, viscosity, solvent properties and pH. Maeda et al., for example, showed that temperature and solvent viscosity affect radical-pair lifetimes and MFE magnitude in E. coli DNA photolyase [9]. Solvent properties can also modify radical-pair formation and recombination [15]. Changes in pH additionally affect flavin protonation equilibria and electron-transfer kinetics, with consequences for radical-pair photochemistry and magnetic-field responses [16,17]. MFE magnitude and response kinetics may therefore carry different information about the environment surrounding a protein-based sensor.

Here, we investigate MagLOV2, an engineered LOV2-domain protein exhibiting magnetic-field-dependent fluorescence, as a genetically programmable sensor for organelle-specific measurements in living cells [4,18].

We examine its magnetic response following localisation to the cytoplasm, nucleus and mitochondria, and compare mitochondrial targeting sequences to determine whether fluorescence intensity can be increased without substantially changing MFE contrast (Fig. 1a,b).

We then follow mitochondrially targeted MagLOV2 during CCCP (carbonyl cyanide m-chlorophenyl hydrazone)-induced mitophagy, in which damaged mitochondria are delivered to acidic lysosomal compartments through the PINK1–Parkin pathway (Fig. 1c) [19–22]. Lysosomal markers and the low-pH mt-Keima signal [23] are used to examine the redistribution of MagLOV2-labelled material, while magnetic-field measurements determine whether its response kinetics change during this process. Measurements of purified protein across a controlled pH range provide a test of the contribution from acidification (Fig. 1d).

Finally, we examine the possible photochemical basis of the pH dependence using structural analysis of the FMN-binding environment, absorption spectroscopy and protonation-dependent kinetic simulations. The analysis considers FMN and nearby residues, including E116 and H117, as potential contributors to the observed kinetics, without assigning the response to a unique radical-pair pathway. In this way, genetic localisation and magnetic-response measurements are used to investigate organelle transport and its associated physicochemical changes through an observable complementary to conventional fluorescence imaging [24].

## RESULTS

### Subcellular localization and magnetic-field response of MagLOV2

To examine whether MagLOV2 retains magnetic-field sensitivity after genetic targeting, we expressed the protein in the cytoplasm, nucleus and mitochondria of HEK293T cells (Fig. 2a–c). Cell boundaries were identified using CellMask Red; nuclear localisation was directed by a simian virus 40 (SV40)-derived nuclear localisation signal and verified by counterstaining; and mitochondrial localisation was achieved using a cytochrome c oxidase subunit 8A (COX8A) targeting sequence and confirmed with MitoTracker Red.

**Fig. 2.**
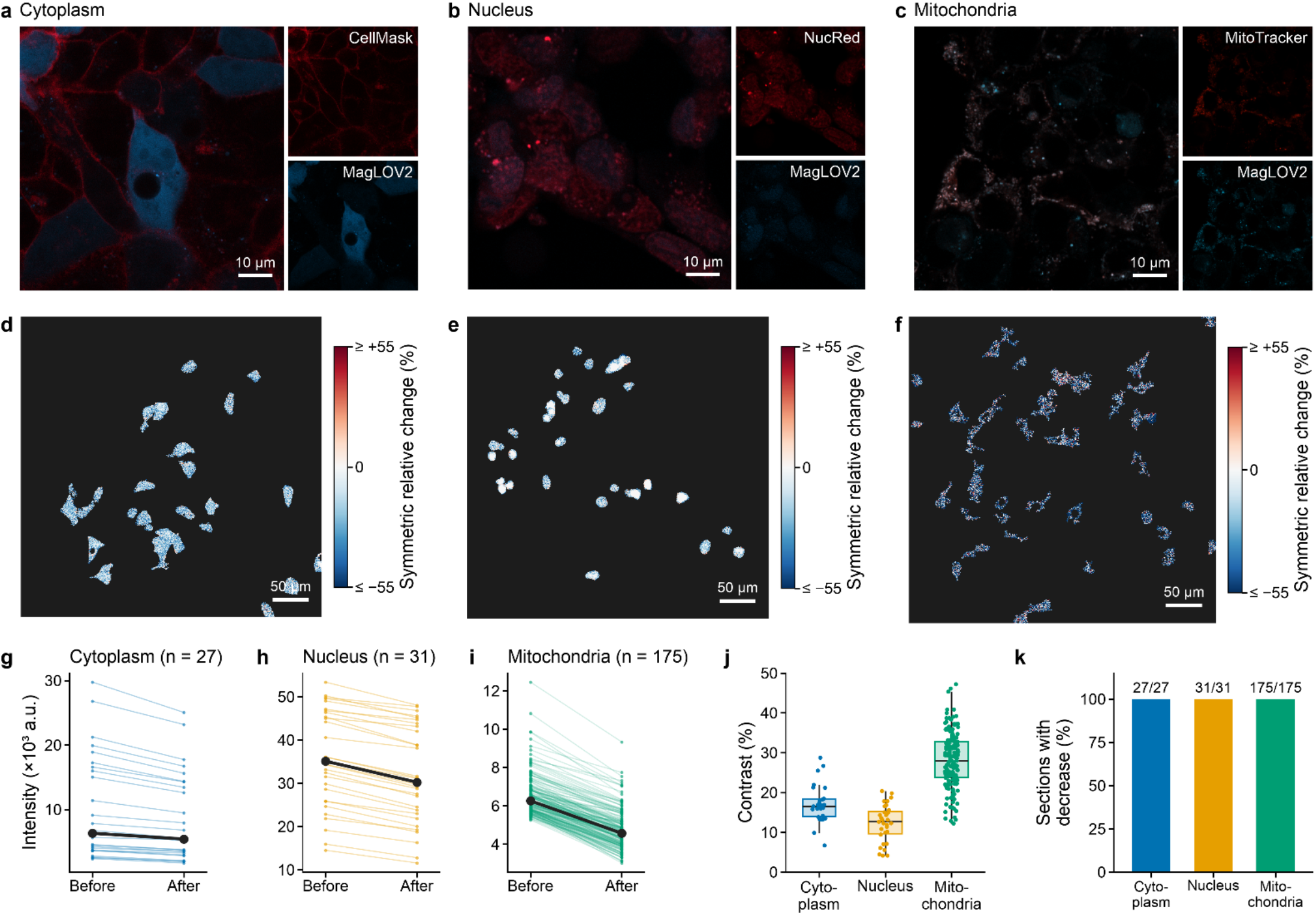
Subcellular targeting and magnetic-field-dependent fluorescence of MagLOV2 in living cells. **a–c**, Fluorescence images of HEK293T cells expressing MagLOV2 in the cytoplasm (a), nucleus (b), and mitochondria (c). CellMask Red was used to visualize cell boundaries, a nuclear counterstain was used to identify nuclei, and MitoTracker Red was used to identify mitochondria. Nuclear and mitochondrial localization was achieved using an SV40-derived nuclear localization signal and COX8A targeting sequence, respectively. **d–f**, Spatial maps of the symmetric relative fluorescence change following application of a 16 mT magnetic field in the cytoplasm (d), nucleus (e), and mitochondria (f). Negative values (blue) indicate decreased fluorescence under the applied magnetic field. **g–i**, Paired fluorescence intensities measured before and after magnetic-field application in cytoplasmic regions (g, n = 27), nuclear regions (h, n = 31), and mitochondrial sections (i, n = 175). Thin lines represent individual regions or sections, and black lines indicate the mean. **j**, Distribution of MFE fluorescence contrast in the three compartments, with mean values of 17.1 ± 4.8%, 12.3 ± 4.7%, and 28.0 ± 6.9% (mean ± SD) for the cytoplasm, nucleus, and mitochondria, respectively. **k**, Percentage of analysed regions exhibiting decreased fluorescence following magnetic-field application. A decrease was observed in all measured regions (27/27 cytoplasmic, 31/31 nuclear, and 175/175 mitochondrial sections).

Application of a 16 mT magnetic field decreased MagLOV2 fluorescence in all three compartments, as shown by the spatial maps and paired intensity measurements (Fig. 2d–i). The mean MFE contrast was 17.1 ± 4.8% in the cytoplasm (mean ± SD, n = 27), 12.3 ± 4.7% in the nucleus (n = 31) and 28.0 ± 6.9% in mitochondria (n = 175). Every analysed region showed a fluorescence decrease: 27/27 cytoplasmic regions, 31/31 nuclear regions and 175/175 mitochondrial sections (Fig. 2k). Mitochondrial MagLOV2 exhibited both the largest mean contrast and a broader distribution of values (Fig. 2j).

The contrast differences may reflect compartment-dependent physicochemical conditions. Mitochondrial microviscosity, for example, could alter radical-pair lifetimes and spin relaxation [9,25,26], although contributions from local protein environment, expression or other compartment-specific factors cannot be excluded. The measurements demonstrate magnetic responsiveness in each of the three targeted locations; they do not establish the origin of the larger mitochondrial contrast.

### Optimization of mitochondrial targeting enhances MagLOV2 fluorescence while preserving magnetic-field sensitivity

We compared N-terminal targeting sequences derived from cytochrome c oxidase subunit 8A (COX8A) and oxidase assembly 1-like protein (OXA1L) to improve the fluorescence signal of mitochondrially localised MagLOV2 [27]. Experiments were performed in HeLa cells, whose elongated morphology facilitated visualisation of mitochondrial structures. Each sequence was fused to MagLOV2 or monomeric ultra-stable GFP (muGFP), allowing the effect of targeting sequence to be compared between fluorescent proteins.

OXA1L-targeted MagLOV2 showed stronger mitochondrial fluorescence than the COX8A-targeted protein, whereas muGFP fluorescence did not differ significantly between targeting sequences (Fig. 3a). Immunofluorescence microscopy using an anti-His antibody, MitoTracker Red and DAPI showed that both MagLOV2 constructs overlapped with the mitochondrial marker (Fig. 3b,c; equivalent muGFP experiments in Fig. S1). The fluorescence enhancement was therefore observed without an apparent loss of mitochondrial localisation and was not reproduced by muGFP.

**Fig. 3.**
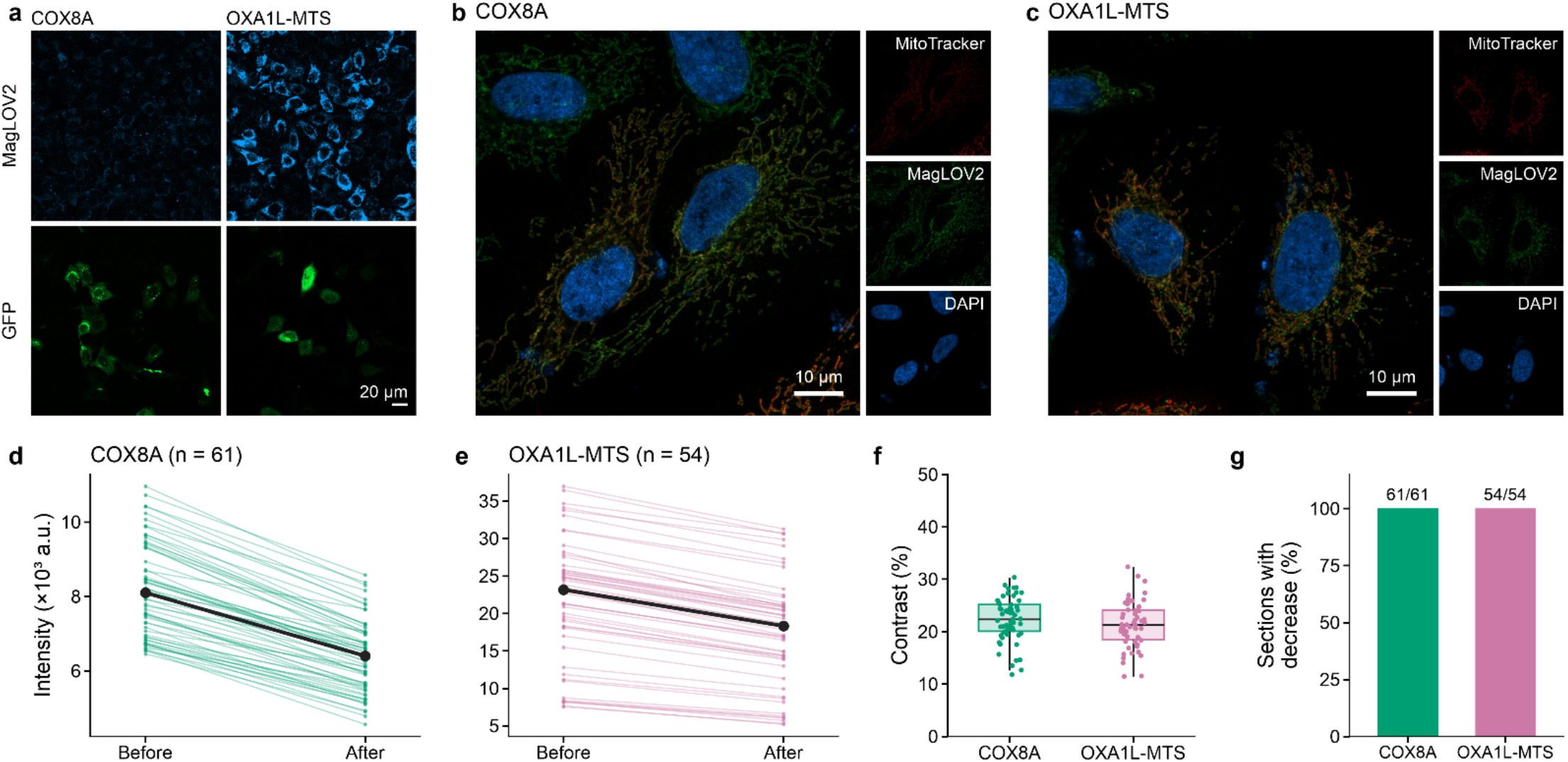
Optimization of mitochondrial targeting improves MagLOV2 fluorescence while preserving magnetic-field responsiveness. **a**, Representative fluorescence images of HeLa cells expressing MagLOV2 (cyan, upper panels) or monomeric ultra-stable GFP (muGFP; green, lower panels) fused to mitochondrial targeting sequences derived from COX8A or OXA1L (OXA1L-MTS). OXA1L-MTS increased MagLOV2 fluorescence relative to COX8A targeting, whereas muGFP fluorescence was comparable between the two constructs. **b**,**c**, Immunofluorescence images of HeLa cells expressing COX8A-targeted (b) or OXA1L-MTS-targeted (c) MagLOV2. MitoTracker Red (red) labels mitochondria, anti-His immunostaining (green) detects MagLOV2, and DAPI (blue) labels nuclei. Both constructs show colocalization with the mitochondrial marker. **d**,**e**, Paired fluorescence intensities measured before and after application of a 16 mT magnetic field in mitochondrial sections expressing COX8A-targeted (d, n = 61) or OXA1L-MTS-targeted (e, n = 54) MagLOV2. Thin lines represent individual mitochondrial sections, and black lines with filled circles indicate the mean. **f**, Distribution of MFE fluorescence contrast for COX8A-targeted and OXA1L-MTS-targeted MagLOV2, with mean contrasts of 22.3 ± 4.2% and 21.3 ± 4.4%, respectively (mean ± SD). **g**, Percentage of mitochondrial sections exhibiting decreased fluorescence following magnetic-field application. Fluorescence decreased in all analysed sections for both constructs (61/61 for COX8A and 54/54 for OXA1L-MTS).

Both constructs remained sensitive to a 16 mT magnetic field, which decreased fluorescence in all analysed mitochondrial sections (Fig. 3d,e,g). Before field application, mean fluorescence intensities were approximately 8.2 × 10^3^ a.u. for COX8A-targeted MagLOV2 and 2.18 × 10^4^ a.u. for the OXA1L-targeted construct, a 2.7-fold difference. The corresponding MFE contrasts were similar: 22.3 ± 4.2% and 21.3 ± 4.4%, respectively (mean ± SD; Fig. 3f). Fluorescence decreased in 61/61 COX8A-targeted and 54/54 OXA1L-targeted mitochondrial sections (Fig. 3g). OXA1L targeting therefore provided a stronger fluorescence signal for subsequent measurements without appreciably changing the relative magnetic-field response.

### Mitophagy-associated acidification alters MagLOV2 magnetic-field response kinetics

We examined MagLOV2 during CCCP-induced mitophagy to determine whether its magnetic-field response changes as labelled mitochondrial material enters a different intracellular environment. HeLa cells constitutively expressing Parkin were treated with the mitochondrial uncoupler CCCP. In the PINK1–Parkin pathway, depolarised mitochondria recruit Parkin and are subsequently delivered to lysosomes for degradation. Mitochondrially targeted MagLOV2 was therefore expected to accompany the damaged mitochondrial material into acidic compartments.

Acidification was assessed using mt-Keima, which has excitation maxima near 440 and 590 nm under neutral and acidic conditions, respectively, and an emission maximum near 620 nm [23]. Following CCCP treatment, the low-pH excitation signals increased while the high-pH signal decreased (Fig. 4a–d). The acidic-to-neutral excitation ratio increased from 1.16 to 6.31, a 5.4-fold change. Although mt-Keima has a reported pKa of approximately 6.5, absolute pH assignment requires calibration of the relative excitation efficiencies. We therefore use this ratio as evidence of increased exposure to acidic compartments, without assigning an exact intracellular pH.

**Fig. 4.**
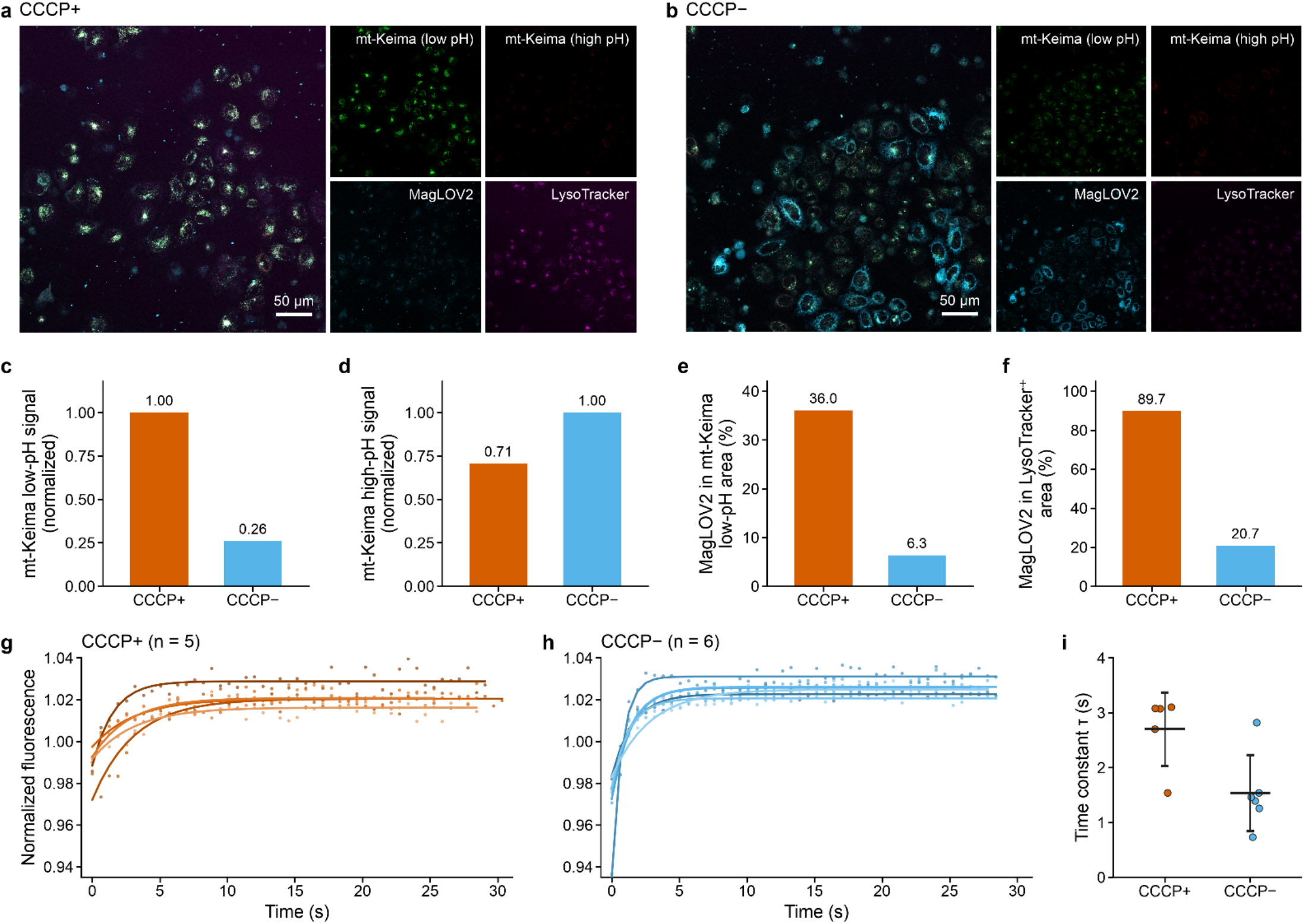
Mitophagy-associated acidification and changes in the magnetic-field response kinetics of MagLOV2. **a**,**b**, Representative fluorescence images of Parkin-expressing HeLa cells expressing mitochondrially targeted MagLOV2 and mt-Keima following CCCP treatment (CCCP+, a) or under untreated conditions (CCCP–,b). Separate channels show mt-Keima signals under low-pH and high-pH excitation conditions, MagLOV2 fluorescence, and LysoTracker staining. **c**,**d**, Normalized mt-Keima fluorescence signals under low-pH (c) and high-pH (d) excitation conditions in CCCP-treated and untreated cells. Values were normalized to the higher signal in each panel. The acidic-to-neutral excitation ratio increased from 1.16 in untreated cells to 6.31 following CCCP treatment. **e**, Percentage of the low-pH mt-Keima-positive area overlapping with MagLOV2 fluorescence in CCCP-treated (36.05%) and untreated (6.34%) cells. **f**, Percentage of LysoTracker-positive area overlapping with MagLOV2 fluorescence in CCCP-treated (89.73%) and untreated (20.75%) cells. **g**,**h**, Normalized magnetic-field-dependent fluorescence responses of MagLOV2 in CCCP-treated (g, n = 5) and untreated (h, n = 6) measurement regions. Each region represents an averaged response from multiple cells within a separate field of view. Points show measured fluorescence values, and solid curves represent single-exponential fits used to determine the response time constant, τ. **i**, Distribution of fitted MFE response time constants in CCCP-treated and untreated regions. Points represent individual measurement regions, with horizontal bars and error bars indicating the mean and standard deviation, respectively. CCCP treatment was associated with slower MFE response kinetics.

The spatial association of MagLOV2 with these compartments was examined using mt-Keima and LysoTracker. The overlap between MagLOV2 and the low-pH mt-Keima signal increased from 6.34% in untreated cells to 36.05% following CCCP treatment (Fig. 4e); overlap with LysoTracker rose from 20.75% to 89.73% (Fig. 4f). These measurements are consistent with delivery of MagLOV2-labelled mitochondrial material to acidic lysosomal compartments during mitophagy.

We next recorded MagLOV2 fluorescence following magnetic-field application in untreated and CCCP-treated cells. Each measurement was an averaged response from multiple cells within a separate field of view. Single-exponential fits yielded the MFE response time constant, τ (Fig. 4g,h). The mean time constant increased from approximately 1.5 s in untreated cells to 2.7 s following CCCP treatment (Fig. 4i). Thus, slower magnetic-response kinetics accompanied the redistribution of MagLOV2 towards acidic compartments. Acidification is a possible contributor, although changes in other local environmental factors during mitophagy cannot be excluded. We examined this dependence further using purified protein under controlled pH conditions.

### pH-dependent magnetic-field response kinetics and MagLOV2 radical-pair photochemistry

To test whether acidity could contribute to the slower response measured during mitophagy, we measured magnetic-field-dependent fluorescence kinetics of purified MagLOV2 in buffers from pH 4.5 to 9.5. Single-exponential fits to the normalised responses gave time constants of 8.27 ± 0.37 s at pH 4.5, 4.77 ± 0.16 s at pH 5.5, 2.60 ± 0.08 s at pH 6.5, 2.61 ± 0.11 s at pH 7.5, 2.53 ± 0.08 s at pH 8.5 and 1.56 ± 0.08 s at pH 9.5 (Fig. 5d,e). The response changed most strongly below pH 6.5, with comparatively little variation between pH 6.5 and 8.5. Its slowing under acidic conditions is consistent with the trend observed in CCCP-treated cells.

**Fig. 5.**
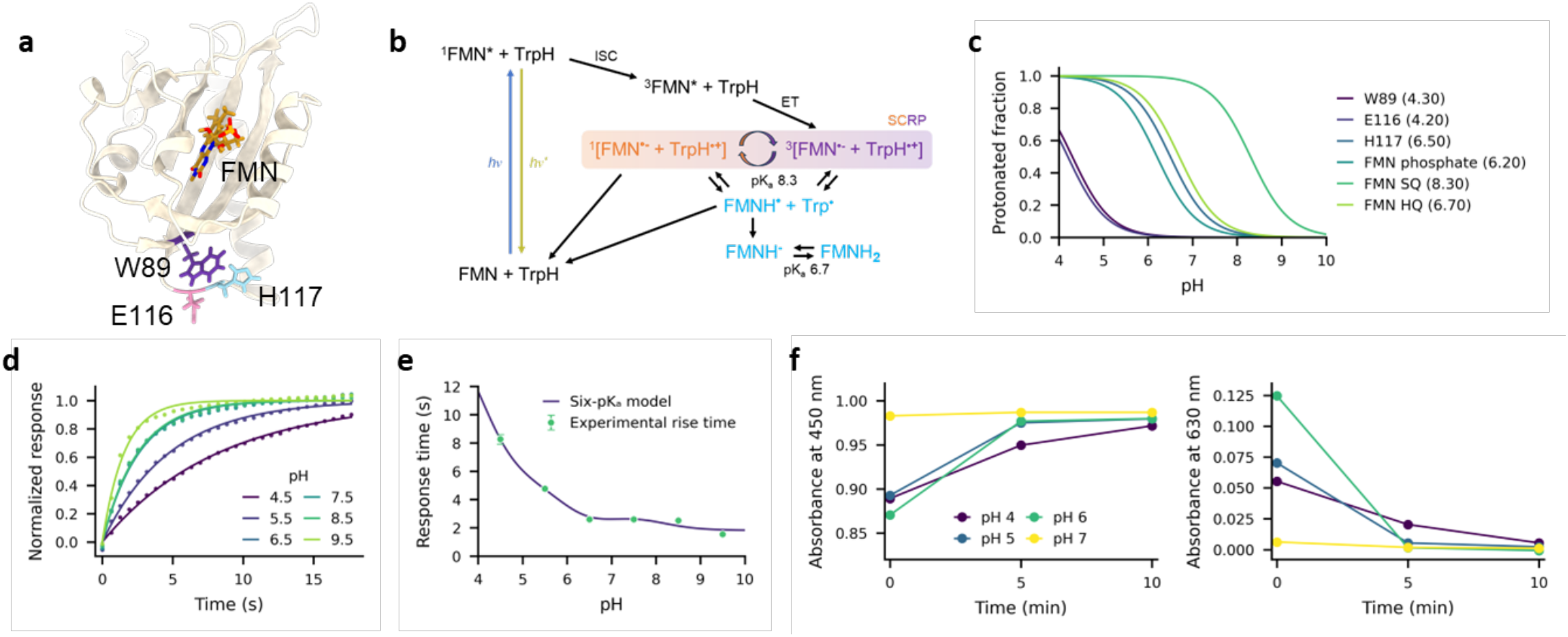
pH-dependent magnetic-field response kinetics of MagLOV2 and the proposed FMN–W89 radical-pair mechanism. **a**, Structural representation of MagLOV2 showing the FMN cofactor and W89, the proposed electron donor in the spin-correlated radical pair. E116 and H117 are located near W89 and may influence radical-pair photochemistry through changes in their protonation states. **b**, Proposed reaction scheme describing photoinduced FMN–W89 radical-pair formation, spin-dependent recombination, and subsequent flavin redox reactions. **c**, Calculated protonated fractions of W89 (pKa = 4.30), E116 (4.20), H117 (6.50), FMN phosphate (6.20), FMN semiquinone (8.30), and FMN hydroquinone (6.70) as a function of pH. The curves were calculated using the Henderson–Hasselbalch equation and assigned pKa values. **d**, Normalized magnetic-field-dependent fluorescence responses of purified MagLOV2 measured at pH 4.5, 5.5, 6.5, 7.5, 8.5, and 9.5, with solid curves showing single-exponential fits. **e**, Experimental MFE response time constants (points) and the simulated pH dependence (purple curve) obtained from a kinetic model incorporating all six protonation equilibria shown in (c). Error bars represent the reported uncertainties in the fitted experimental time constants. **f**, Time-dependent normalized absorbance of MagLOV2 at 450 and 630 nm at pH 4, 5, 6, and 7, measured at 0, 5, and 10 min. The spectral changes are consistent with pH-dependent flavin redox-state interconversion during illumination.

We examined the FMN-binding environment and flavin-centred reaction network to identify possible sources of this pH dependence (Fig. 5a,b). Photoexcited FMN can generate spin-correlated radical intermediates through electron transfer, and their formation, spin evolution and recombination can influence the magnetic-field-dependent fluorescence response. Previous work on engineered MagLOV proteins has identified flavin– protein geometry, electron-transfer kinetics and spin relaxation as determinants of magnetic sensitivity [18]; studies of cryptochromes have also shown the sensitivity of flavin radical-pair chemistry to the protein environment [12]. In MagLOV2, W89 lies close to FMN and may participate in the electron-transfer network, whereas E116 and H117 may affect the nearby electrostatic and protonation environment. These assignments identify possible contributions without requiring a single defined radical-pair pathway.

We considered six acid–base equilibria: W89, E116, H117, FMN phosphate, FMN semiquinone and FMN hydroquinone (Fig. 5c). Their different pH dependences provide possible means of modifying electron transfer, proton exchange, radical-state interconversion, recombination and flavin recovery. To investigate their combined effect, we developed a kinetic model that describes interconversion between oxidised FMN, neutral and anionic semiquinone, and hydroquinone states. Effective reaction rates depend on the specified protonation equilibria, with exchange between semiquinone states and protonation-dependent hydroquinone oxidation included explicitly (Fig. 5b). The simulated response times reproduced the principal experimental trends: slower response under acidic conditions, nearly constant times from pH 6.5 to 8.5 and a faster response at pH 9.5 (Fig. 5e). Among the alternatives examined, scenarios containing multiple local protonation equilibria described the experimental behaviour better than scenarios dominated by a single protonation event (Fig. S2). This agreement supports a contribution from several protonation-dependent processes but does not uniquely identify their individual rates or molecular pathways.

We also recorded UV–visible absorption spectra from pH 4 to 7. Initial spectra showed oxidised-FMN bands between 400 and 500 nm, including prominent absorption near 450 and 475 nm. During illumination, absorption near 450 nm increased and absorption near 630 nm decreased at pH 4–6; the changes were substantially smaller at pH 7 (Fig. 5f; Fig. S3). Oxidised FMN contributes strongly near 450 nm, whereas the longer-wavelength band is consistent with flavin radical absorption. The opposing changes therefore indicate redistribution among flavin photochemical states in a pH-dependent manner.

To examine this redistribution, we decomposed spectra measured at 0, 5 and 10 min into oxidised FMN, semiquinone and hydroquinone contributions using a reversible photochemical model (Fig. S4-5). The reconstructed spectra reproduced the principal absorption features and their evolution during illumination. The fitted populations likewise differed with pH, consistent with changes in flavin-state interconversion. Together with the kinetic calculations, these spectroscopic measurements support the interpretation that protonation-sensitive flavin photochemistry contributes to the pH dependence of MagLOV2 response kinetics. The roles of W89, E116 and H117 remain proposed contributions, and the data do not resolve a unique sequence of electron-transfer or radical-pair reactions.

## DISCUSSION

Our results demonstrate that a genetically encoded quantum sensor can follow a defined biological structure as it moves between intracellular compartments while remaining sensitive to changes in its local physicochemical environment. MagLOV2 was successfully targeted to the cytoplasm, nucleus, and mitochondria of living mammalian cells, retaining magnetic-field-dependent fluorescence in each compartment. During CCCP-induced mitophagy, mitochondrially localised MagLOV2 was delivered to acidic lysosomal compartments, accompanied by a slower magnetic-field response. Measurements of purified MagLOV2 subsequently established a pronounced dependence of the response time on pH, while structural analysis, absorption spectroscopy, and kinetic simulations allowed the possible molecular origins of this behaviour to be examined. These observations extend the application of protein-based quantum sensing to the investigation of physicochemical changes associated with intracellular transport and organelle degradation.

A principal advantage of genetically encoded quantum sensors is their ability to remain associated with a specific biological component during intracellular transport. Conventional quantum sensors, including nitrogen-vacancy centres in nanodiamonds and molecular quantum nanosensors, have enabled measurements of physicochemical parameters within living cells, but generally require external delivery and are not intrinsically associated with genetically specified cellular structures [1,2]. MagLOV2, by comparison, can be expressed directly within a target compartment and subsequently transported with the labelled material. Following CCCP treatment, the overlap between mitochondrially targeted MagLOV2 and LysoTracker increased from 20.75% to 89.73%, while colocalisation with the low-pH mt-Keima signal increased from 6.34% to 36.05%. The corresponding increase in the acidic-to-neutral mt-Keima excitation ratio, from 1.16 to 6.31, independently indicated that the labelled mitochondrial material had entered an acidic environment. Genetic localisation therefore allows the magnetic response to be associated with a biological structure throughout its transport between compartments, including the accompanying changes in its physicochemical surroundings.

The slower MFE response observed during mitophagy is consistent with the pH dependence measured using purified MagLOV2. Under controlled buffer conditions, the response time decreased from 8.27 ± 0.37 s at pH 4.5 to 4.77 ± 0.16 s at pH 5.5, remained approximately 2.6 s between pH 6.5 and 8.5, and reached 1.56 ± 0.08 s at pH 9.5. The pronounced slowing under acidic conditions is particularly relevant to mitophagy, during which mitochondrial material is transported from its original environment into acidic lysosomal compartments. Whereas conventional fluorescence intensity and excitation-ratio measurements provide optical observables of the local environment, the pH dependence of MagLOV2 is also reflected in the temporal response to an applied magnetic field. MFE contrast and response kinetics can therefore be measured independently using the same protein, with the latter providing additional information about changes in the intracellular environment, providing a route toward intracellular multiparameter quantum sensing.

The molecular origin of this pH dependence can be examined through the flavin-centred photochemistry of MagLOV2 and the surrounding protein environment. Photoexcitation of FMN can initiate electron-transfer reactions that generate spin-correlated radical intermediates, whose formation, interconversion, and recombination influence the magnetic-field-dependent fluorescence response [8–13,17,28–30]. Previous investigations of engineered MagLOV proteins and other flavin-based radical-pair systems have shown that magnetic sensitivity depends on electron-transfer geometry, radical recombination, spin relaxation, and interactions with the local protein environment [12,18]. In MagLOV2, W89 lies near the FMN chromophore and may participate in the electron-transfer network, while the neighbouring residues E116 and H117 could influence the local electrostatic and protonation environment. Their proximity to the flavin provides possible routes through which changes in pH could affect the photochemical reaction kinetics, although the contributions of individual residues and radical-pair pathways remain unresolved.

The kinetic simulations suggest that the observed pH dependence may arise from the combined influence of several protonation equilibria within the flavin-centred photochemical network. The model incorporated W89, E116, H117, the FMN phosphate group, and the semiquinone and hydroquinone states of FMN. Because these groups undergo protonation changes over different pH ranges, changes in the surrounding environment may affect multiple reaction steps, including electron transfer, proton exchange, radical-state interconversion, recombination, and flavin recovery. The simulated response times reproduced the main experimental trends, including slower responses under acidic conditions, comparatively little variation between pH 6.5 and 8.5, and faster responses at pH 9.5. The agreement is consistent with a mechanism in which the measured response reflects the combined kinetics of several protonation-dependent processes. It does not, however, uniquely establish the contribution of any individual residue or reaction pathway.

The absorption measurements provide further information about the pH-dependent flavin photochemistry. During illumination, the spectra showed changes in absorption near 450 and 630 nm, with a more pronounced response under acidic conditions. The 450-nm region is dominated by oxidised FMN, whereas absorption near 630 nm is consistent with contributions from flavin radical states, including neutral FMN semiquinone. At pH 4–6, the increase in absorption near 450 nm was accompanied by a decrease near 630 nm, while substantially smaller changes occurred at pH 7. These opposing changes indicate a redistribution of flavin photochemical states during illumination, with the extent of this redistribution depending on pH. Spectral decomposition into oxidised, semiquinone, and hydroquinone contributions reproduced the principal absorption features and indicated differences in the populations and interconversion of these states.

Changes in flavin protonation provide a possible explanation for the spectral behaviour and its relationship to MFE response kinetics. Both the semiquinone and hydroquinone states undergo protonation equilibria within the investigated pH range, allowing environmental pH to influence the relative populations and interconversion rates of oxidised, radical, and reduced flavin species. Under acidic conditions, protonation of reduced flavin states may alter electron-transfer and proton-transfer kinetics, together with subsequent flavin reoxidation. Such changes could affect the accumulation and decay of semiquinone intermediates and prolong the time required for the photochemical system to recover towards its oxidised ground state. Slower recovery of radical or reduced flavin populations is therefore a plausible contribution to the longer magnetic-field response times measured under acidic conditions, although the absorption measurements alone do not establish the individual kinetic steps responsible.

The combined spectroscopic and kinetic results are consistent with pH-dependent changes in flavin redox-state populations and their interconversion influencing the temporal magnetic response of MagLOV2. The protonation-dependent simulations reproduce the experimental response-time behaviour, while the absorption measurements independently show that flavin photochemistry changes with pH. Neither analysis requires the magnetic response to be assigned to a single predetermined radical-pair pathway. Instead, the observed kinetics may reflect the behaviour of a network of flavin-centred electron-transfer and photochemical reactions, influenced by the protonation state of the chromophore and its surroundings. W89, E116, and H117 remain plausible contributors to this behaviour through their effects on local electron transfer, electrostatics, and protonation, but their individual mechanistic roles have yet to be determined experimentally.

In addition to the pH-dependent response kinetics, MagLOV2 exhibited different MFE contrasts following targeting to distinct intracellular compartments. The mean contrast was largest in mitochondria (28.0%), compared with 17.1% in the cytoplasm and 12.3% in the nucleus. These differences may arise, at least in part, from variations in the local physicochemical environment. Radical-pair magnetic responses are sensitive to temperature, viscosity, and solvent conditions, which can affect radical lifetimes, molecular motion, and spin relaxation [9,14]. Previous studies have identified differences in microviscosity among intracellular compartments, including comparatively viscous and heterogeneous mitochondrial environments [25,26]. Such conditions could influence the radical-pair dynamics of MagLOV2 and contribute to the larger mitochondrial contrast. However, the relationship between mitochondrial microviscosity and the measured MFE has not been established directly, and contributions from other compartment-specific conditions cannot be excluded.

The choice of mitochondrial targeting sequence also affected the fluorescence performance of MagLOV2. Replacing the COX8A targeting sequence with OXA1L-MTS increased mitochondrial fluorescence approximately 2.7-fold, while the mean MFE contrast remained similar between the two constructs (22.3 ± 4.2% and 21.3 ± 4.4%, respectively). A comparable fluorescence enhancement was not observed for muGFP, suggesting that the effect depends on properties of MagLOV2 or its interaction with the targeting construct, rather than a general influence of the targeting sequence on fluorescent proteins. The molecular basis of this difference remains unclear. Nevertheless, the increased fluorescence was obtained without an appreciable change in relative magnetic-field sensitivity, indicating that the targeting sequence can influence signal intensity independently of MFE contrast. This is particularly useful for measurements in confined or weakly fluorescent intracellular regions, where the number of detected photons limits the precision of temporal fluorescence measurements.

Genetically encoded quantum-sensitive proteins may also be applicable to biological processes beyond the mitophagy model examined here. Fluorescent proteins have become widely used tools for following specific proteins, organelles, and cellular processes in living systems. Magnetic-field-sensitive photochemistry introduces a further measurable property that may respond to changes in the local environment independently of conventional fluorescence intensity. The behaviour observed for MagLOV2 suggests possible applications to organelle maturation, intracellular trafficking, autophagy, metabolic responses, and physicochemical changes associated with cellular stress. Improvements in sensitivity, calibration, and compatibility with established imaging methods would be necessary before extending such measurements to complex physiological conditions. These developments may be relevant to studies of mitochondrial dysfunction, lysosomal disorders, neurodegeneration, and other conditions involving altered organelle homeostasis. More generally, measurements of spin-dependent photochemistry within living cells may help establish how quantum processes behave in biological environments.

The present work demonstrates that a protein-based quantum sensor can be genetically targeted to specific intracellular compartments and remain associated with labelled material during organelle transport. In MagLOV2, magnetic-field-dependent fluorescence provides a measure of MFE contrast, while the response kinetics exhibit a pronounced dependence on pH. The intracellular and purified-protein measurements, supported by absorption spectroscopy and kinetic simulations, suggest that protonation-dependent flavin photochemistry contributes to the slower response observed during mitophagy-associated acidification. Quantitative interpretation in living cells will require further calibration and a more detailed understanding of the relevant reaction pathways and environmental contributions. Nevertheless, the ability to follow a defined biological structure while measuring changes in magnetic-response kinetics provides an experimental basis for investigating spin-dependent photochemical processes during intracellular biological activity.

## MATERIALS & METHODS

### Plasmid construction and subcellular targeting of MagLOV2

MagLOV2 was genetically fused to targeting sequences designed to direct the sensor to the nucleus or mitochondria. Nuclear localization was achieved using an SV40-derived nuclear localization sequence. For mitochondrial localization, targeting modules derived from human COX8A and OXA1L were fused to the N terminus of MagLOV2. These constructs were used to evaluate organelle-specific localization and to compare the performance of different mitochondrial targeting strategies.

### Protein expression and purification

C-terminally His-tagged MagLOV2 was expressed in E. coli using a pET21b-based construct and a DE3 expression strain. Protein expression was induced with IPTG at reduced temperature, after which cells were harvested and stored until purification. Cell pellets were resuspended in Tris-based lysis buffer, disrupted by sonication, and clarified by centrifugation. The soluble fraction was purified by nickel-affinity chromatography followed by size-exclusion chromatography. The final protein was exchanged into a HEPES-based buffer containing NaCl, EDTA, and DTT and concentrated to approximately 2 mg/mL. MagLOV2 concentration was determined from absorbance at 280 nm using an extinction coefficient of 38,340 M^−1^ cm^−1^ and a molecular mass of 20,236.4 Da.

### Immunofluorescence staining and confocal imaging

Cells cultured on glass-bottom dishes were fixed with paraformaldehyde, permeabilized with Triton X-100, and blocked with bovine serum albumin. MagLOV2 and mitochondria were immunostained using antibodies against the His tag and TOM20, respectively, followed by fluorescently labelled secondary antibodies. Nuclei were counterstained with Hoechst 33342. Samples were mounted in an antifade medium and imaged using a confocal microscope.

### Experimental response times and protonation descriptors

MagLOV2 response times were analyzed at pH 4.5, 5.5, 6.5, 7.5, 8.5 and 9.5. The corresponding measured time constants were 8.274, 4.772, 2.595, 2.614, 2.530 and 1.555 s. Six candidate acid–base equilibria were represented by the Henderson–Hasselbalch protonated fraction

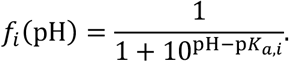

The assigned p*K*_*a*_values were W89, 4.30; E116, 4.20; H117, 6.50; FMN phosphate, 6.20; FMN semiquinone, 8.30; and FMN hydroquinone, 6.70. These values were held fixed during parameter fitting.

### Chemical kinetics

The reduced model tracked neutral FMN semiquinone (*H*, FMNH^•^), anionic semiquinone (*A*, FMN^•−^), and a lumped hydroquinone pool (*Q*). The oxidized fraction was defined by conservation, *O* = 1 –*H* – *A* – *Q*. The governing equations were

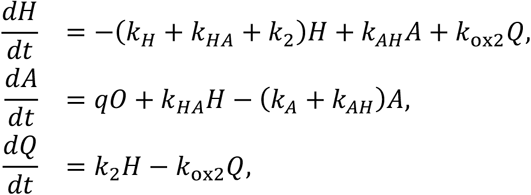

where *q* is an effective photochemical source rate. With **x** = (*H, A, Q*)^⊤^, the equations can be written *d***x**/*dt* = ***M*x** + **b**, with

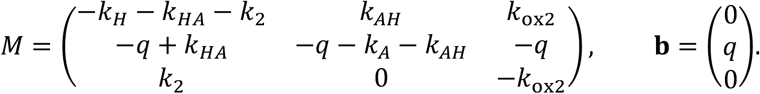

The neutral and anionic semiquinone recovery rates were related by *k*_H_= 0.4*k*_)_. For each fitted scenario, the anionic recovery rate was parameterized as

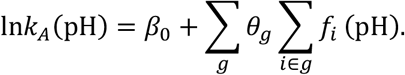

A group *g* contained either one site or sites constrained to share a single coefficient *θ*_*g*_. The semiquinone protonation equilibrium (p*K*_*a*_= 8.30) entered through

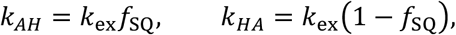

where *A*→*H* is protonation. Hydroquinone oxidation was represented by

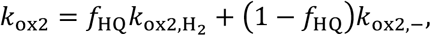

using the hydroquinone p*K*_*a*_ of 6.70. Thus, the semiquinone and hydroquinone p*K*_*a*_values entered every kinetic scenario, regardless of which additional residue or phosphate terms were fitted.

The default constants were *q* = 0.022 s^−1^, *k*_2_ = 0.012 s^−1^, 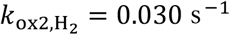, *k*_ox2,−_ = 0.100 s^−1^, and *k*_ex_ = 1.0 s^−1^.

### Response-time calculation

At each pH, steady states were calculated for source rates *q* and *q*_off_ = 0.95*q*:

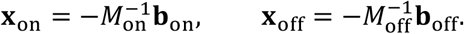

For *Δ***x** = **x**_off_ − **x**_on_, the integral relaxation time of the neutral semiquinone response was calculated as

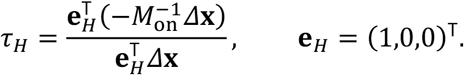

This integral time was compared with the experimental time constants.

### Parameter estimation and model comparison

Recovery parameters were estimated with bounded nonlinear least squares by minimizing the difference between predicted and measured response times with a ridge penalty on the protonation coefficients:

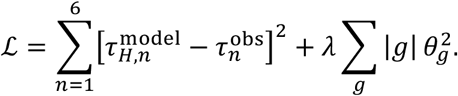

The penalty coefficient was λ = 0.035 and the factor |*g*| counted sites represented by a common coefficient. Bounds on β_0_ were [−8,3] ; coefficient bounds were [−8,8]. Thirteen predefined residue/phosphate grouping scenarios were compared. The best scenario assigned separate fitted coefficients to W89, E116, H117 and FMN phosphate, while the two flavin acid–base equilibria remained fixed as described above.

Each scenario was also assessed by leave-one-out cross-validation (LOOCV). Each of the six pH measurements was withheld in turn, the parameters were refitted using the remaining five measurements, and the omitted time constant was predicted. The root-mean-square error of the six held-out predictions was used to rank the scenarios. The separate individual-descriptor ranking applied the same fitting and LOOCV procedure to six single-descriptor recovery laws; in those additional tests, the two flavin p*K*_*a*_values continued to act through their fixed kinetic terms as well.

### Absorption spectroscopy and spectral preprocessing

UV–visible absorption spectra were analyzed at pH 4, 5, 6 and 7. Measurements acquired at 0, 5 and 10 min were used to estimate FMN photoreduction kinetics. Spectra were interpolated to a common wavelength grid spanning 400–650 nm in 1-nm increments. A baseline estimated as the median absorbance between 680 and 700 nm was subtracted. The corrected spectra were normalized to the absorbance at 450 nm of the corresponding initial reference spectrum. A dark-state spectrum recorded before irradiation was used where available; otherwise, the 0-min spectrum served as the reference.

### Spectral decomposition

The spectra were modelled as linear combinations of oxidized FMN (FMNox), neutral semiquinone (FMNH^•^) and hydroquinone (FMNH^−^). The oxidized-FMN reference was taken from the pH-matched dark spectrum and smoothed with a Gaussian filter of 1-nm standard deviation. Semiquinone and hydroquinone reference spectra were reconstructed from Gaussian components fitted to the corresponding standards. The calculated absorbance was

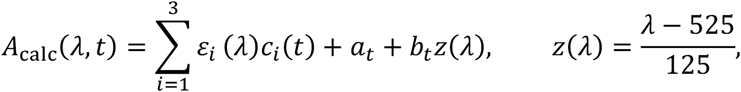

where ε_*i*_(λ) is the reference spectrum, *c*_*i*_(*t*) the fractional population of species *i*, and *a*_C_ and *b*_C_ the scan-specific offset and linear-baseline corrections. Populations were constrained to be nonnegative and to sum to unity.

### Photoreduction kinetics

Photoreduction was represented by two consecutive reversible steps:

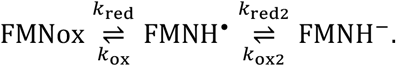

The fractional populations *c*_ox_, *c*_sQ_ and *c*_HQ_ obeyed

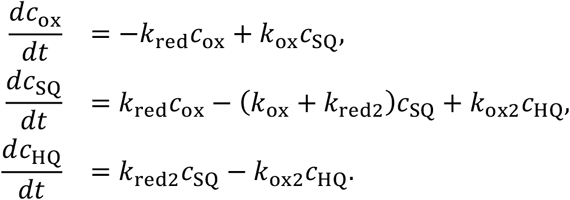

Writing **c** = (c_ox_, *c*_sQ_, *c*_HQ_)^⊤^ gives

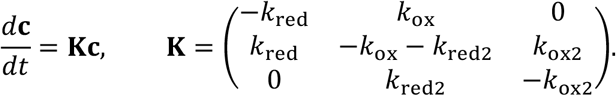

Populations were evaluated by matrix exponentiation, **c**(*t*) = exp(**K***t*)**c**(0). Time was expressed in minutes, and kinetic rate constants therefore had units of min^−1^.

### Parameter estimation

Initial species fractions and four rate constants were fitted jointly for each pH condition. Initial fractions were constrained to be positive and normalized to sum to one. Rate constants were optimized in logarithmic coordinates. The penalized objective combined spectral residuals with regularization of log rates and scan-specific baseline terms:

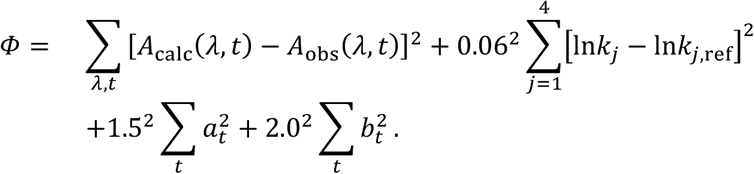

Reference rates were 0.04, 0.04, 0.02 and 0.02 min^−1^ for *k*_red_, *k*_ox_, *k*_red2,_ and *k*_ox2_, respectively. Rates were bounded between 10^−^ and 5 min^−1^, offsets to ±0.08 normalized absorbance units and baseline slopes to ±0.06. The objective was minimized using nonlinear least squares. Fits using an external oxidized-FMN reference and independently estimated populations at each time point were also evaluated as spectral reconstruction controls.

### Fit assessment

Spectral residuals were summarized by the root-mean-square error,

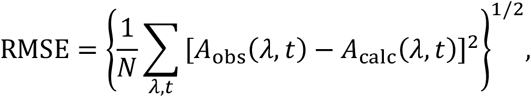

where *N* is the number of spectral observations. Approximate local uncertainties were estimated from the Jacobian of the regularized least-squares objective and used for plotting; they were not interpreted as confidence intervals. Three irradiation time points per pH limited independent identification of the four rate constants, particularly where reference spectra overlapped.

## REFERENCES

1. Yukawa, H. et al. Quantum life science: biological nano quantum sensors, quantum technology-based hyperpolarized MRI/NMR, quantum biology, and quantum biotechnology. Chem. Soc. Rev. 54, 3293–3322 (2025).

2. Ishiwata, H., Song, J., Shigeno, Y., Nishimura, K. & Yanai, N. Molecular quantum nanosensors functioning in living cells. Sci. Adv. 12, eaeb5422 (2026).

3. Jones, N. Why ‘quantum proteins’ could be the next big thing in biology. Nature 651, 25–27 (2026).

4. Abrahams, G. et al. Quantum spin resonance in engineered proteins for multimodal sensing. Nature 649, 1172–1179 (2026).

5. Burd, S. C. et al. Magnetic resonance control of spin-correlated radical pair dynamics in vivo. Nature 651, 940–945 (2026).

6. Feder, J. S. et al. A fluorescent-protein spin qubit. Nature 645, 73–79 (2025).

7. Meng, K. et al. Optically detected and radio wave-controlled spin chemistry in flavoproteins. Nat. Biotechnol. 10.1038/s41587-026-03158-5 (2026).

8. Henbest, K. B. et al. Magnetic-field effect on the photoactivation reaction of Escherichia coli DNA photolyase. Proc. Natl Acad. Sci. USA 105, 14395–14399 (2008).

9. Maeda, K. et al. Magnetically sensitive light-induced reactions in cryptochrome are consistent with its proposed role as a magnetoreceptor. Proc. Natl Acad. Sci. USA 109, 4774–4779 (2012).

10. Evans, E. W., Dodson, C. A., Maeda, K., Biskup, T., Wedge, C. J. & Timmel, C. R. Magnetic field effects in flavoproteins and related systems. Interface Focus 3, 20130037 (2013).

11. Sheppard, D. M. W. et al. Millitesla magnetic field effects on the photocycle of an animal cryptochrome. Sci. Rep. 7, 42228 (2017).

12. Gravell, J. et al. Spectroscopic characterization of radical pair photochemistry in nonmigratory avian cryptochromes: magnetic field effects in GgCry4a. J. Am. Chem. Soc. 147, 24286–24298 (2025).

13. Antill, L. M., Kohmura, M., Jimbo, C. & Maeda, K. Introduction of magneto-fluorescence fluctuation microspectroscopy for investigating quantum effects in biology. Nat. Photonics 19, 178–186 (2025).

14. Steiner, U. E. & Ulrich, T. Magnetic field effects in chemical kinetics and related phenomena. Chem. Rev. 89, 51–147 (1989).

15. Hoang, H. M., Pham, V. T. B., Grampp, G. & Kattnig, D. R. Magnetic field-sensitive radical pair dynamics in polymethylene ether-bridged donor–acceptor systems. ACS Omega 3, 10296–10305 (2018).

16. Murakami, M., Maeda, K. & Arai, T. Dynamics of intramolecular electron transfer reaction of FAD studied by magnetic field effects on transient absorption spectra. J. Phys. Chem. A 109, 5793–5800 (2005).

17. Antill, L. M. & Woodward, J. R. Flavin adenine dinucleotide photochemistry is magnetic field sensitive at physiological pH. J. Phys. Chem. Lett. 9, 2691–2696 (2018).

18. Antill, L. M., Baidoo, J. K. & Gerhards, L. Revealing essential properties for enhanced quantum sensing in engineered AsLOV2 proteins. Preprint at bioRxiv 10.64898/2026.02.27.708212 (2026).

19. Geisler, S. et al. PINK1/Parkin-mediated mitophagy is dependent on VDAC1 and p62/SQSTM1. Nat. Cell Biol. 12, 119–131 (2010).

20. Lazarou, M. et al. The ubiquitin kinase PINK1 recruits autophagy receptors to induce mitophagy. Nature 524, 309–314 (2015).

21. Gladkova, C., Maslen, S. L., Skehel, J. M. & Komander, D. Mechanism of parkin activation by PINK1. Nature 559, 410–414 (2018).

22. Zhu, Q. et al. GRAF1 integrates PINK1-Parkin signaling and actin dynamics to mediate cardiac mitochondrial homeostasis. Nat. Commun. 14, 8187 (2023).

23. Sun, N. et al. A fluorescence-based imaging method to measure in vitro and in vivo mitophagy using mt-Keima. Nat. Protoc. 12, 1576–1587 (2017).

24. Chudakov, D. M., Matz, M. V., Lukyanov, S. & Lukyanov, K. A. Fluorescent proteins and their applications in imaging living cells and tissues. Physiol. Rev. 90, 1103–1163 (2010).

25. Chambers, J. E. et al. An optical technique for mapping microviscosity dynamics in cellular organelles. ACS Nano 12, 4398–4407 (2018).

26. Pytowski, L. et al. Mapping viscosity in discrete subcellular locations with a BODIPY based fluorescent probe. Preprint at bioRxiv 10.1101/522532 (2019).

27. Stiburek, L. et al. Knockdown of human Oxa1l impairs the biogenesis of F1Fo-ATP synthase and NADH oxidoreductase. J. Mol. Biol. 374, 506–516 (2007).

28. Antill, L. M., Takizawa, S.-Y., Murata, S. & Woodward, J. R. Photoinduced flavin-tryptophan electron transfer across vesicle membranes generates magnetic field sensitive radical pairs. Mol. Phys. 117, 2594–2603 (2019).

29. Antill, L. M., Beardmore, J. P. & Woodward, J. R. Time-resolved optical absorption microspectroscopy of magnetic field sensitive flavin photochemistry. Rev. Sci. Instrum. 89, 023707 (2018).

30. Antill, L. M. & Vatai, E. RadicalPy: a tool for spin dynamics simulations. J. Chem. Theory Comput. 20, 9488–9499 (2024).

