## Supplementary material for "Organelle-Specific pH Quantification Using a Protein-Based Quantum Sensor": SI

### TITLE

Underline: first author(s), *: corresponding author(s)

ORCID: HI: 0000-0002-7486-8772; LMA: 0000-0001-9266-5584;

### Contents

S1: Mitochondrial localization of muGFP fusion constructs in HeLa cells S3

S2: Comparison of $pK_a$-dependent kinetic models S4

S3: Time-dependent changes in MagLOV2 absorption spectra S5

S4: Reference spectra and calculated flavin populations S6

S5: Comparison of measured and calculated absorption spectra S7

#### S1: Mitochondrial localization of muGFP fusion constructs in HeLa cells

The intracellular distribution of muGFP fusion constructs containing different mitochondrial targeting sequences (MTS) was examined by fluorescence microscopy. Constructs incorporating sequences from COX8A and OXA1L were compared with mitochondrial staining to assess their subcellular distribution.

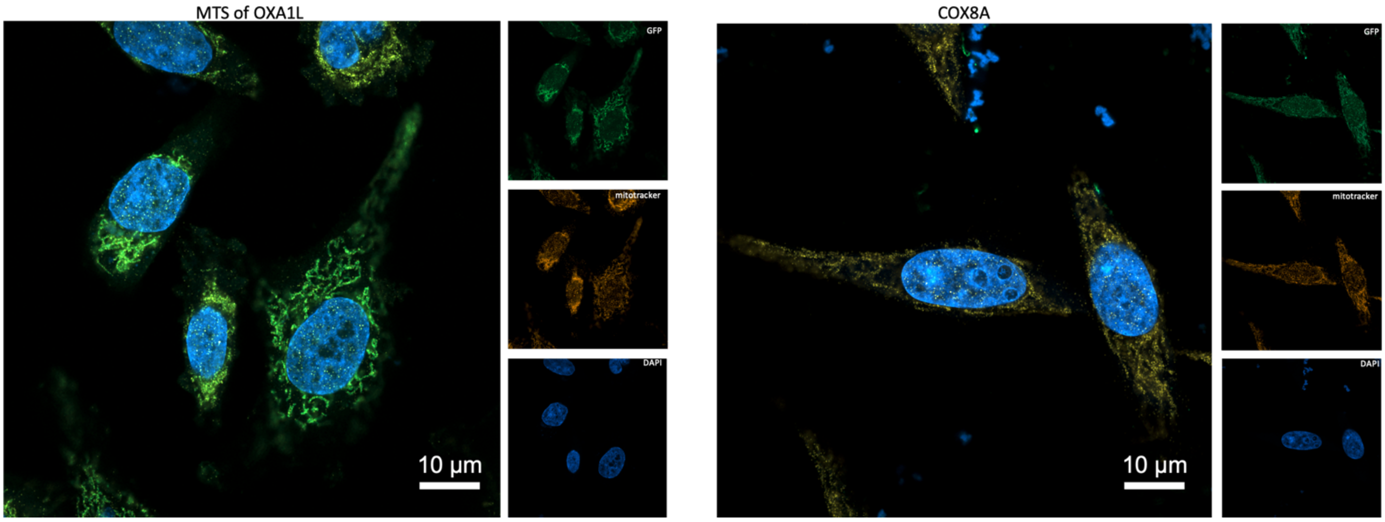


**Figure S1 | Localization of muGFP fusion constructs containing different mitochondrial targeting sequences.**

Representative fluorescence images of HeLa cells expressing muGFP fusion constructs containing MTS derived from COX8A and OXA1L. muGFP fluorescence is shown in green, mitochondrial staining in orange and nuclear staining in blue. Merged images and the corresponding individual fluorescence channels are shown for each construct.

#### S2: Comparison of $pK_a$-dependent kinetic models

The pH dependence of the measured MFE response times was compared with kinetic models containing different combinations of candidate protonation-dependent terms assigned to components of the local flavin environment. Predictive performance was assessed by leave-one-out cross-validation (LOOCV), in which each pH measurement was excluded in turn and predicted using parameters fitted to the remaining measurements. These model terms were used to test alternative protonation-dependent descriptions of the kinetics and do not, by themselves, establish a unique electron-transfer or radical-pair pathway.

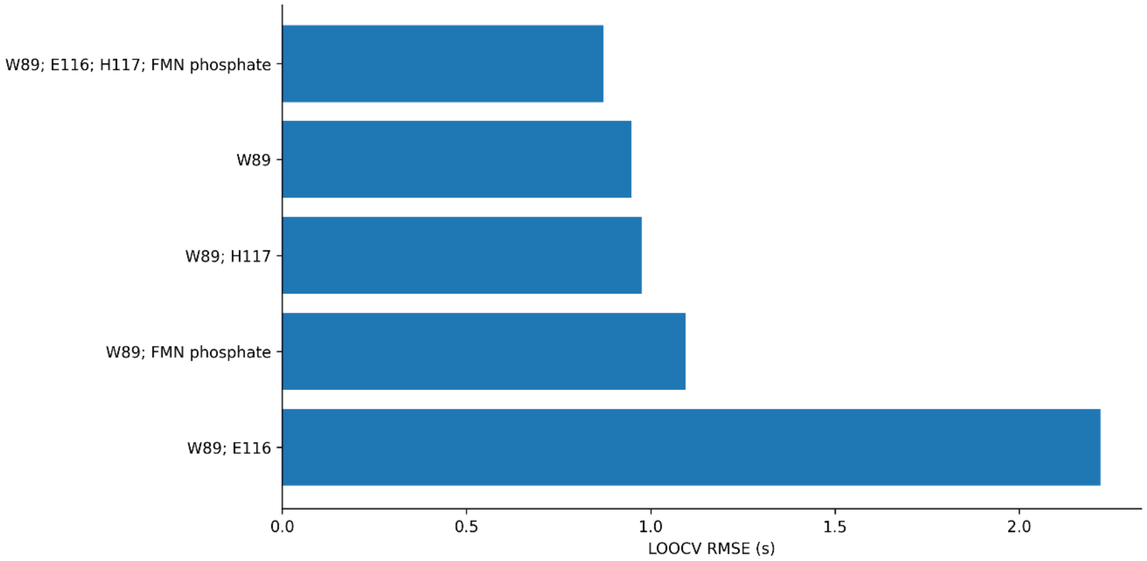


**Figure S2 | Leave-one-out cross-validation of $pK_a$-dependent kinetic models.**

Root-mean-square error (RMSE) of the predicted MFE response time for five kinetic models incorporating different combinations of candidate protonation-dependent terms. The models include a W89-associated term alone, W89-associated and H117 terms, W89-associated and FMN-phosphate terms, W89-associated and E116 terms, and a model in which W89-associated, E116, H117, and FMN-phosphate terms contribute independently. A lower RMSE indicates better agreement between the excluded measurement and its prediction. The FMN semiquinone and hydroquinone acid-base equilibria are included in the underlying kinetic scheme for all five models. The residue labels denote candidate protonation terms used for model comparison and should not be interpreted as identifying a unique electron donor or radical partner.

#### S3: Time-dependent changes in MagLOV2 absorption spectra

Changes in the absorption spectra of purified MagLOV2 were followed during illumination at pH 4–7. Spectra recorded at 0, 5, and 10 min were compared with the corresponding dark-state reference where available. Absorbance at 450 and 630 nm was extracted to examine changes in the oxidized-flavin absorption region and the longer-wavelength region associated with flavin radical-state contributions.

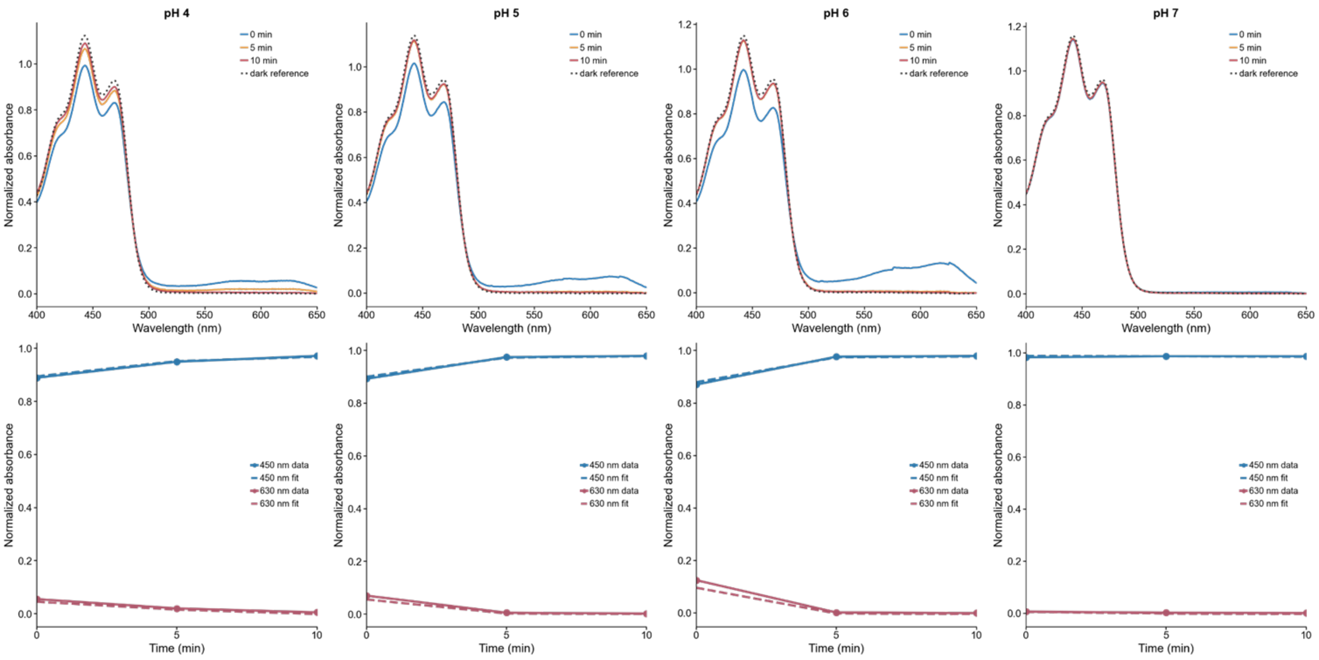


**Figure S3 | Time-dependent absorption spectra of MagLOV2 at different pH values.**

Upper panels: normalized absorption spectra measured at pH 4, 5, 6 and 7 after 0, 5 and 10 min of illumination. Dark-state reference spectra are also shown. Lower panels: normalized absorbance at 450 and 630 nm as a function of illumination time. Symbols represent the measured values and dashed lines show the corresponding values calculated from the fitted spectral model. All spectra were baseline-corrected and normalized using the initial absorbance at 450 nm.

#### S4: Reference spectra and calculated flavin populations

The measured absorption spectra were analyzed using reference spectra representing oxidized FMN, neutral flavin semiquinone, and a reduced flavin hydroquinone state. The resulting kinetic model was used to calculate the time-dependent contribution of each spectral component at pH 4–7. These values are model-derived fractional populations and should not be interpreted as independently measured concentrations of individual flavin species.

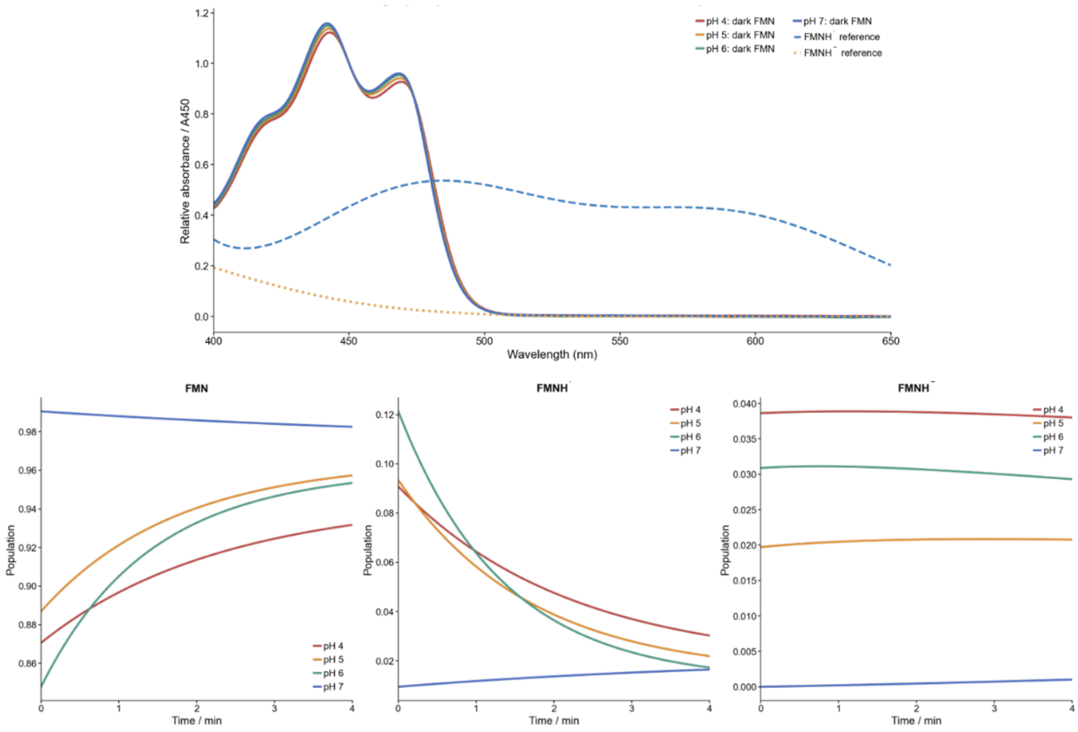


**Figure S4 | Reference spectra and calculated populations of flavin redox states.**

Upper panel: reference absorption spectra used for spectral decomposition. Oxidized-FMN reference spectra are shown for the four pH conditions, together with reference spectra assigned to neutral semiquinone (FMNH•) and a reduced flavin state represented as $FMNH^-$ in the three-state spectral model. Lower panels: calculated model-derived fractional populations of oxidized FMN, FMNH•, and the reduced flavin component during the first 4 min of illumination at pH 4–7. Population profiles were obtained from the fitted reversible three-state kinetic model and represent the components required to reproduce the measured spectra rather than direct measurements of chemical-species concentrations.

#### S5: Comparison of measured and calculated absorption spectra

The agreement between the measured absorption spectra and the fitted kinetic model was examined across the full wavelength and illumination-time ranges. Residual maps were calculated by subtracting the model-reconstructed absorbance from the experimental absorbance at each wavelength and time point.

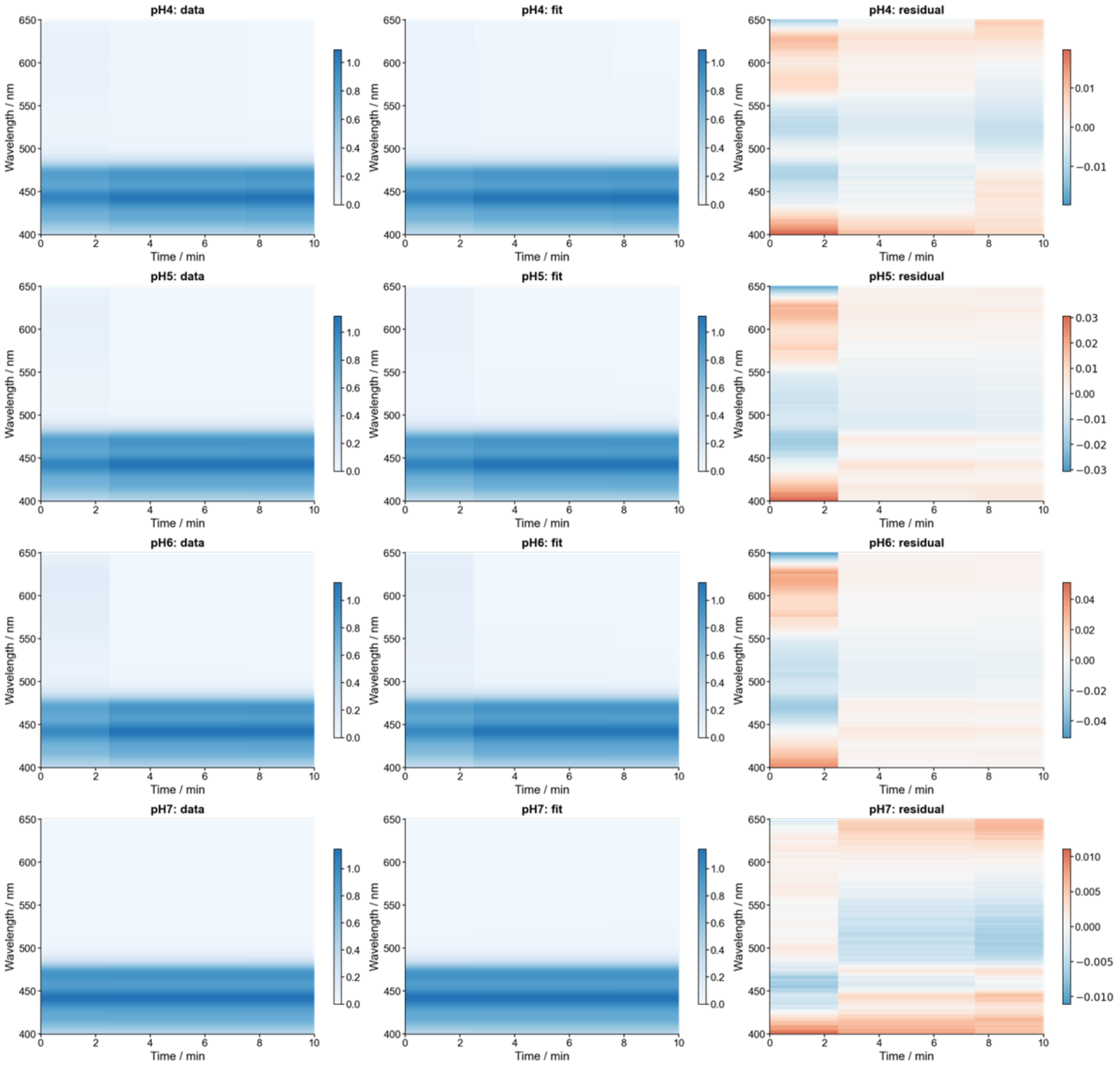


**Figure S5 | Measured absorption spectra, kinetic fits and residuals at pH 4–7.**

Two-dimensional maps of normalized absorbance as a function of wavelength and illumination time. The left column shows the measured spectra, the middle column shows spectra reconstructed from the fitted kinetic model, and the right column shows the residuals (measured minus model-reconstructed absorbance). Rows correspond to pH 4, 5, 6 and 7. The spectral range is 400–650 nm, and the measurements span 0–10 min of illumination. Residual color scales indicate deviations in normalized absorbance units.
